# Whole-Blood Transcriptomics Highlights Galectin-1 in Systolic Blood Pressure

**DOI:** 10.64898/2026.09.11.751025

**Authors:** Amadou Gaye, Ashton F. Oliver, Merry L. Lindsey, German E. Gonzalez

## Abstract

**Background:** Galectins regulate inflammation, fibrosis, immunity, and vascular remodeling. Although galectin-3 has been widely studied in cardiovascular disease, the relative contributions of other galectins to blood pressure regulation remain unclear. We systematically evaluated associations between galectin gene expression and systolic blood pressure (SBP) in African Americans.

**Methods:** Whole-blood mRNA sequencing data were analyzed from 419 African American participants in the GENE-FORECAST cohort who were free of type 2 diabetes and not receiving antihypertensive medications. LGALS1, LGALS3, LGALS8, and LGALS9 were evaluated using generalized linear models adjusted for age, sex, and body mass index. Associations with cardiac magnetic resonance imaging traits and circulating inflammatory cytokines were also examined.

**Results:** LGALS1 showed the strongest association with SBP (β=13.25 mm□Hg per standard deviation increase; P=7.57×10□^1^□). LGALS9 was positively associated with SBP (β=9.73; P=7.59×10□□), whereas LGALS3 showed a weaker positive association (β=4.06; P=0.019). LGALS8 was inversely associated with SBP (β=−6.45; P=1.66×10□□), as was the LGALS3/LGALS1 ratio (β=−10.2; P=2.34×10□□). Galectin genes also showed distinct associations with cardiac imaging traits. Cytokine associations were modest and inconsistent, although LGALS1 was positively associated with interleukin-1β.

**Conclusions:** Whole-blood transcriptomics identified LGALS1 as the strongest galectin correlate of SBP in African Americans. Divergent associations across galectin genes suggest heterogeneous, potentially nonredundant cardiovascular roles. These findings broaden the focus beyond LGALS3 and support further investigation of LGALS1 as a biomarker and possible mediator of hypertension-related vascular stress.

## INTRODUCTION

Hypertension remains the leading modifiable contributor to cardiovascular morbidity and mortality worldwide and disproportionately affects African Americans, who exhibit earlier disease onset, higher prevalence, and greater risk of hypertensive complications than other populations ^1-3^. Despite substantial advances in antihypertensive therapy, the molecular pathways underlying inter-individual variability in blood pressure regulation and cardiovascular remodeling remain incompletely understood ^4,5^. Identification of transcriptional biomarkers reflecting vascular stress, inflammation, and tissue remodeling may improve mechanistic understanding and support precision medicine approaches to hypertension.

Galectins are a family of β-galactoside-binding proteins involved in immune regulation, extracellular matrix remodeling, fibrosis, and vascular biology ^6-8^. Among them, LGALS3 (Galectin 3) has received substantial attention in cardiovascular research because of its established associations with cardiac fibrosis, inflammation, heart failure, and vascular remodeling ^9-11^. Elevated circulating galectin-3 levels have been associated with adverse cardiovascular outcomes and hypertensive target-organ damage ^11-13^. In contrast, other galectin family members, including LGALS1 (Galectin 1), LGALS8 (Galectin 8), and LGALS9 (Galectin 9), remain comparatively understudied in the context of blood pressure regulation ^1,13^. Importantly, most prior studies have focused on individual galectins in isolation rather than systematically evaluating the relative contribution of multiple galectin family members within the same population ^14-17^.

Emerging evidence suggests that galectins may exert distinct and potentially complementary functions in cardiovascular homeostasis. Experimental studies implicate LGALS1 in endothelial function, immune modulation, angiogenesis, and vascular smooth muscle biology ^18, 19^. Galectin-1 has also been proposed to participate in compensatory anti-inflammatory and tissue-remodeling responses under cardiovascular stress conditions ^20-22^. Meanwhile, LGALS8 and LGALS9 have been linked to inflammatory signaling, immune-cell regulation, cardiometabolic phenotypes, and vascular dysfunction ^23^. However, whether these galectin family members exhibit differential transcriptional relationships with systolic blood pressure (SBP) in humans remains unclear. Furthermore, the relative contribution of individual galectin family members to blood pressure-associated cardiovascular phenotypes has not been systematically characterized in African Americans, a population disproportionately affected by hypertension and hypertensive complications ^24,25^.

In the present study, we systematically evaluated the whole-blood expression profiles of galectin family members in relation to SBP and cardiovascular phenotypes in African Americans from the Genomics, Environmental Factors and the Social Determinants of Cardiovascular Disease in Africans Americans (GENE-FORECAST) Study cohort. To minimize candidate-driven bias, all galectin genes with sufficient expression levels were analyzed using identical statistical models. We further examined associations with cardiac imaging traits, inflammatory cytokines, and circulating cardiovascular protein biomarkers. Through this comparative transcriptomic framework, we identified the galectin family members most strongly associated with SBP and cardiovascular phenotypes in untreated African Americans.

## MATERIALS AND METHODS

### Study Cohort

GENE-FORECAST is a research platform strategically designed to employ a comprehensive, multi-omics systems biology approach for in-depth, multi-dimensional phenotyping of health and disease within the African American population. Utilizing a community-based sampling framework, GENE-FORECAST established a cohort comprising 669 U.S.-born African American men and women aged 21-65, recruited primarily from the metropolitan Washington D.C. area as described previously ^26^. The GENE-FORECAST Study was approved by the Institutional Review Boards of the National Institutes of Health and Meharry Medical College. All participants provided written informed consent prior to their involvement. The project presented here was conducted in compliance with local regulations and institutional protocols.

### RNA Sequencing Data

GENE-FORECAST whole□blood gene expression profiles were obtained through messenger RNA sequencing (mRNA□seq). Total RNA was extracted from blood samples preserved in stabilization tubes using the MagMAX™ RNA isolation kit (Life Technologies, Carlsbad, CA). Library preparation was performed from total RNA using Illumina TruSeq reagents, producing indexed complementary DNA (cDNA) libraries following ribosomal RNA depletion.

Paired□end RNA sequencing was conducted using Illumina HiSeq 2500 and HiSeq 4000 platforms, with each sample sequenced to a depth of at least 50 million reads. Gene expression levels were quantified using the Broad Institute’s GTEx RNA□seq processing workflow, following their publicly available analysis pipeline^2^. Transcripts exhibiting low expression were filtered out prior to normalization, defined as those with fewer than 2 counts per million (CPM) detected in at least three samples. Normalization of expression data was performed using the Trimmed Mean of M□values (TMM) method, which is appropriate for count□based RNA□sequencing data^3^. Principal component analysis was subsequently applied to identify expression outliers, resulting in the removal of four transcripts from downstream analyses. A total of 17,947 protein□coding passed quality□control procedures. Four galectin genes (LGALS1, LGALS3, LGALS8, and LGALS9) demonstrated sufficient expression for robust statistical analysis.

### Statistical Analyses

Analyses were restricted to GENE-FORECAST participants who were not receiving antihypertensive medications. This restriction was applied to minimize pharmacological and metabolic confounding, because blood pressure–lowering therapies may influence both SBP and systemic transcriptional profiles.

Associations between galectin gene expression and SBP were evaluated using generalized linear models, with SBP modeled as the dependent variable and galectin gene expression as the independent variable of interest. All galectin family members with sufficient whole-blood expression levels, including LGALS1, LGALS3, LGALS8, and LGALS9, were evaluated systematically using identical models to minimize candidate-driven bias. Rather than acting in isolation, these carbohydrate-binding proteins operate as an integrated system for tissue homeostasis ^27^. Specifically, LGALS1 and LGALS3 exert opposing, highly coordinated effects on the cardiovascular system, whilst LGALS1 functions primarily as a homeostatic, anti-inflammatory mediator, LGALS3 acts as a potent driver of macrophage activation and severe cardiac fibrosis. Consequently, the LGALS3/LGALS1 expression ratio was also computed and analyzed using the same modeling framework, to understand the dynamic interplay between these pro and anti-inflammatory molecules.

All models were adjusted for age, sex, and body mass index (BMI), given their known relationships with blood pressure variation and systemic gene-expression profiles ^28,29^. To facilitate comparison of effect sizes across galectin family members with differing expression ranges, normalized gene-expression values were standardized to z-scores prior to regression analyses; consequently, beta coefficients represent the mean difference in blood pressure (mmHg) associated with a one-standard-deviation increase in galectin expression. Beta coefficients, 95% confidence intervals, and two-sided P-values were estimated for each association. Statistical significance was defined as P ≤ 0.05. All analyses were performed in R version 4.4.1 (2024-06-14).

## RESULTS

### Study Population Characteristics

The study included 419 African American participants with available whole-blood RNA-sequencing and clinical data. None were receiving antihypertensive medications, minimizing pharmacological confounding. All participants were free of type 2 diabetes. Measurements of estimated Glomerular Filtration Rate (eGFR) were available for all except 4 participants. The cohort exhibited broad variability in blood pressure measurements, with mean systolic blood pressure (SBP) of 147 ± 37 mmHg and approximately half of participants meeting criteria for hypertension. SBP correlated positively with age, BMI, diastolic blood pressure, inflammatory markers, and left ventricular mass, while exhibiting an inverse correlation with estimated glomerular filtration rate (Table 1).

**Table 1:** Baseline demographic, clinical, and cardiometabolic characteristics of the 419 participants included in the analyses. Values are presented as mean ± standard deviation or count (proportion, %), as appropriate. Abbreviations: SBP, systolic blood pressure; DBP, diastolic blood pressure; BMI, body mass index; eGFR, estimated glomerular filtration rate; BP, blood pressure; T2D, type 2 diabetes mellitus.

| Characteristics | N | Mean<br>or<br>Count | Standard<br>Deviation or<br>Proportion (%) | Correlation<br>(r) with SBP | P of Correlation<br>with SBP |
| --- | --- | --- | --- | --- | --- |
| Age | 419 | 45 | 11 | 0.26 | 5.77e-08 |
| Sex | 419 |  |  | -0.17 | 4.00E-04 |
| Female |  | 247 | 58.9% |  |  |
| Male |  | 172 | 41.1% |  |  |
| Hypertensive | 419 |  |  | 0.85 | 1.15e-119 |
| No |  | 208 | 49.6% |  |  |
| Yes |  | 211 | 50.4% |  |  |
| **SBP (mm Hg)** | 419 | 147 | 37 | - | - |
| **DBP (mm Hg)** | 419 | 74 | 10 | 0.34 | 4.76e-13 |
| **BMI (kg/m<sup>2</sup>)** | 419 | 30 | 7 | 0.15 | 0.002 |
| **eGFR (mL/min/1.73m<sup>2</sup>)** | 415 | 100 | 20 | -0.28 | 9.09E-09 |
| **BP medication** | 419 |  |  | - | - |
| No |  | 419 | 100.0% |  |  |
| Yes |  | 0 | 0.0% |  |  |
| **T2D** | 419 |  |  | - | - |
| No |  | 419 | 100.0% |  |  |
| Yes |  | 0 | 0.0% |  |  |

### Galectin Genes Expression Landscape in the Whole-Blood mRNA-seq Data

Among galectin family members detected in whole blood, four genes demonstrated expression levels sufficient for robust statistical analysis (**Figure 1**): LGALS1, LGALS3, LGALS8, and LGALS9. All four genes were systematically evaluated using identical generalized linear models adjusted for age, sex, and BMI. In addition, the LGALS3/LGALS1 expression ratio was analyzed to assess potential balance between the two most biologically characterized galectins.

**Figure 1.**
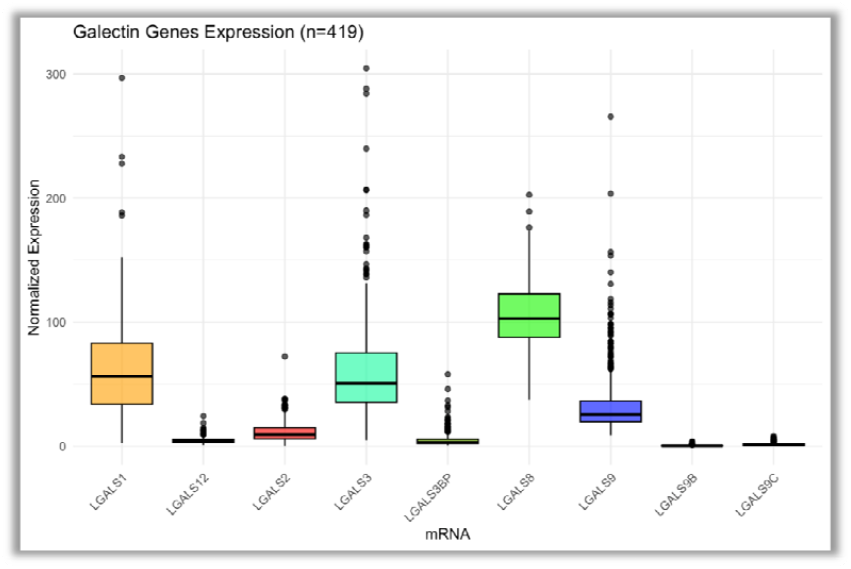
Whole-blood expression landscape of galectin family genes in captured in the mRNA-sequencing data. Boxplots show normalized mRNA expression levels for galectin family members detected in the RNA-sequencing dataset. Among the evaluated genes, LGALS1, LGALS3, LGALS8, and LGALS9 demonstrated sufficient expression levels and passed QC filters for downstream statistical analyses. Central lines represent medians, boxes indicate interquartile ranges, and whiskers denote 1.5 × interquartile range. Outliers are shown as individual points.

### Association Between Galectin Gene Expression and Systolic Blood Pressure

Among all evaluated galectin family members, LGALS1 demonstrated the strongest association with SBP (β = 13.25, P = 7.57 × 10^-15^), substantially exceeding the magnitude and statistical significance observed for other galectin genes (**Table 2**). LGALS9 also exhibited a strong positive association with SBP (β = 9.73, P = 7.59 × 10^-9^), whereas LGALS3 demonstrated a smaller positive association (β = 4.06, P = 0.019). In contrast, LGALS8 was inversely associated with SBP (β = −6.45, P = 1.66 × 10^-4^). Notably, the LGALS3/LGALS1 ratio was inversely associated with SBP (β = -10.2, P = 2.34 × 10^-9^), suggesting that the relative balance between these galectins may reflect distinct transcriptional states associated with elevated blood pressure. Collectively, these analyses identified LGALS1 as the dominant whole-blood galectin gene associated with SBP in this cohort.

**Table 2:** Results of the associations between standardized galectin gene-expression levels and SBP in 41 African samples. Gene-expression values were standardized to z-scores prior to analysis; therefore, bet coefficients represent the change in SBP (mmHg) associated with a one-standard-deviation change in galectin expression. The LGALS3/LGALS1 ratio was before the standardization for analysis. Confidence intervals (95% CI) and two-sided P-values are shown for each association.

| mRNA | Beta | P | 95%CI |
| --- | --- | --- | --- |
| LGALS1 | 13.25 | $7.57 \times 10^{-15}$ | 10.03, 10.46 |
| LGALS3 | 4.06 | 0.019 | 0.68, 7.44 |
| LGALS8 | -6.45 | $1.66 \times 10^{-4}$ | -9.78, -3.12 |
| LGALS9 | 9.73 | $7.59 \times 10^{-9}$ | 6.5, 12.96 |
| LGALS3/LGALS1 | -10.2 | $2.34 \times 10^{-9}$ | -13.48, -6.93 |

### Associations with Cardiac Structure and Function

To explore whether galectin transcriptional signatures extended beyond SBP, we evaluated associations with cardiac imaging phenotypes derived from cardiac magnetic resonance imaging, including left ventricular end-diastolic volume, end-systolic volume, stroke volume, cardiac output, and ejection fraction, which were available in a subset of participants. The results, summarized in the **Supplemental Material**, showed that LGALS3 demonstrated positive associations with stroke volume (N = 140, β = 4.28, P = 0.01, 95% CI = 1.07, 7.5) and ejection fraction (N = 140, β = 1.80, P = 0.009, 95% CI = 0.48, 3.13), while LGALS9 correlated positively with end-diastolic volume (N = 248, β = 3.66, P = 0.05, 95% CI = -0.01, 7.32), and cardiac output (N = 140, β = 0.39, P = 8.39 × 10^-6^, 95% CI = 0.23, 0.56). In contrast, LGALS8 exhibited a strong inverse association with cardiac output (N = 140, β = −0.49, P = 2.10 × 10^-5^, 95% CI = −0.71, −0.27).

Although LGALS1 associated positively with ejection fraction (N = 140, β = 1.84, P = 0.02, 95% CI = 0.30, 3.39), its most pronounced and consistent relationship remained with SBP rather than cardiac structural measures. These results indicate that distinct galectin family members may preferentially associate with different cardiovascular phenotypes, with LGALS1 showing the clearest direct systemic relationship with blood pressure itself.

### Associations with Circulating Inflammatory Cytokines

We next assessed whether galectin transcriptional profiles were associated with 4 circulating inflammatory cytokines, namely interleukin-1 beta (IL-1β), interleukin-6 (IL-6), interleukin-10 (IL-10), tumor necrosis factor alpha (TNF-α), measured from the serum samples available for 378 individuals. Several galectin genes demonstrated nominal associations with inflammation. LGALS1 correlated positively with IL-1β (N = 378, β = 0.12, P = 0.004, 95% CI = 0.04, 0.21), and the ratio LGALS3/LGALS1 associated positively IL-1β (N = 378, β = −0.11, P = 0.01, 95% CI = −0.19, −0.03). These associations were weaker and more heterogeneous than the robust relationship observed between LGALS1 and SBP. None of the galectin family members demonstrated statistically significant associations consistently across all four evaluated inflammatory cytokines. The complete cytokine association results for all galectin genes are provided in the **Supplemental Material**.

## DISCUSSION

In this study, we evaluated the transcriptional relationship between multiple galectin family members and systolic blood pressure in African Americans using whole-blood mRNA sequencing data. By applying identical analytical models across all sufficiently expressed galectin genes, we identified substantial heterogeneity in the strength and direction of association between individual galectins and SBP. Among all evaluated genes, LGALS1 emerged as the dominant correlate of SBP, exhibiting a markedly stronger association than LGALS3, the galectin family member most extensively studied in cardiovascular disease. In addition, distinct galectin family members demonstrated divergent associations with cardiac imaging phenotypes and circulating inflammatory cytokines, suggesting potentially nonredundant roles in cardiovascular and inflammatory biology. Our findings support the concept that the galectin family represents a transcriptionally heterogeneous network in hypertension rather than a biologically uniform group of biomarkers.

### LGALS1 emerges as the dominant galectin associated with SBP

The most striking finding of this study was the magnitude and statistical robustness of the association between LGALS1 expression and SBP. LGALS3 has historically dominated the cardiovascular galectin literature because of its established links to fibrosis, heart failure, and vascular remodeling ^6,30^. However, our results indicate that LGALS1 may exhibit a stronger transcriptional relationship with systolic blood pressure regulation itself, at least in whole blood from untreated African Americans. Importantly, this observation emerged from a systematic comparison framework in which all galectin genes were evaluated under identical statistical conditions, minimizing candidate-driven bias.

Several biological mechanisms may plausibly explain the strong relationship between LGALS1 and SBP. Galectin-1 has been implicated in endothelial biology, angiogenesis, immune-cell regulation, oxidative stress responses, and vascular smooth muscle cell function ^18, 19^. Experimental studies further suggest that galectin-1 may modulate inflammatory signaling and tissue-remodeling responses during cardiovascular stress conditions ^31^. In vascular disease models, galectin-1 has been proposed to exert compensatory or context-dependent effects that may limit pathological remodeling and inflammation ^21^. The strong positive association observed in our data may therefore reflect activation of systemic vascular stress-response pathways accompanying elevated blood pressure rather than a direct pathogenic effect alone.

Notably, LGALS1 showed its strongest and most consistent relationship with SBP itself rather than with cardiac structural phenotypes. This pattern may indicate that LGALS1 transcription primarily reflects systemic vascular or inflammatory states associated with blood pressure elevation before the emergence of overt cardiac remodeling. Longitudinal studies will be necessary to determine whether elevated LGALS1 expression precedes hypertension progression or instead reflects downstream adaptation to chronic hemodynamic stress.

### Differential cardiovascular signatures across galectin family members

Our findings also demonstrate that individual galectin family members exhibit distinct cardiovascular association profiles. LGALS9 showed a strong positive association with SBP and additional relationships with cardiac output and end-diastolic volume, suggesting possible involvement in inflammatory or hemodynamic adaptation pathways. Galectin-9 is increasingly recognized as an important regulator of immune-cell activation and immune checkpoint signaling ^32^, but its role in hypertension remains poorly characterized. The present findings suggest that LGALS9 warrants further investigation in cardiovascular and vascular-inflammatory contexts.

In contrast, LGALS8 demonstrated inverse associations with SBP and cardiac output, suggesting potentially divergent biological functions compared with LGALS1 and LGALS9. Galectin-8 has previously been linked to immune signaling, endothelial activation, and cardiometabolic phenotypes ^33^, although mechanistic cardiovascular studies remain limited. The inverse directionality observed here raises the possibility that some galectin family members may participate in protective or compensatory pathways during blood pressure dysregulation.

LGALS3 exhibited comparatively modest associations with SBP despite its extensive characterization in cardiovascular disease. However, LGALS3 showed positive relationships with stroke volume and ejection fraction, suggesting that its strongest transcriptional associations in this cohort may relate more closely to myocardial remodeling and cardiac functional adaptation than to blood pressure itself. This interpretation aligns with the broader literature implicating galectin-3 in fibrosis, extracellular matrix remodeling, and heart failure progression rather than primary blood pressure regulation ^6,30^.

### Potential biological relevance of the LGASL3/LGALS1 ratio

An additional notable observation was the association between the LGALS3/LGALS1 expression ratio and SBP. Although individual galectins have often been studied independently, these findings suggest that the relative balance between galectin family members may provide additional biological information beyond isolated gene-expression levels. Because galectin-1 and galectin-3 can exert overlapping yet context-dependent effects on inflammation, fibrosis, immune regulation, and tissue remodeling ^34,35^, the ratio between these molecules may reflect broader shifts in vascular or inflammatory transcriptional states accompanying hypertension. Future mechanistic studies are needed to determine whether this ratio captures coordinated regulatory programs or compensatory interactions within the galectin network.

### Inflammatory associations were more modest than blood pressure associations

Although several galectin genes demonstrated nominal associations with inflammatory cytokines, these relationships were generally weaker and less consistent than those observed with SBP. LGALS1 showed a positive association with IL-1β, supporting prior evidence linking galectin-1 to inflammatory and immune-regulatory pathways ^19,36^. However, none of the evaluated galectin family members demonstrated consistent associations across all cytokines examined. These findings suggest that the transcriptional relationship between galectin genes and blood pressure may not simply reflect generalized systemic inflammation captured by conventional circulating cytokines. Instead, galectin expression may integrate multiple biological processes relevant to vascular homeostasis, immune signaling, endothelial stress, and tissue remodeling.

### Importance of studying hypertension biology in African Americans

A major strength of this study is the focus on African Americans, a population disproportionately affected by hypertension and hypertensive complications yet historically underrepresented in transcriptomic cardiovascular studies ^26,37^. African Americans experience earlier hypertension onset, greater disease severity, and higher rates of hypertensive target-organ damage compared with other populations ^38-40^. Despite these disparities, most molecular studies of hypertension have been conducted predominantly in European ancestry populations. Our findings therefore contribute to addressing an important representation gap in cardiovascular genomics and transcriptomics.

In addition, analyses were restricted to individuals not receiving antihypertensive medications and predominantly free of type 2 diabetes, reducing potential confounding from pharmacological treatment and advanced metabolic disease. This design strengthens the interpretation that observed transcriptional associations are more directly related to blood pressure biology itself rather than secondary treatment effects.

## Conclusions

Systematic transcriptomic profiling of galectin family members identified LGALS1 as the dominant whole-blood correlate of systolic blood pressure in African Americans. Distinct galectin family members demonstrated divergent associations with blood pressure, cardiac phenotypes, and inflammatory markers, supporting the concept that galectins participate in heterogeneous and potentially nonredundant cardiovascular pathways. These findings broaden current understanding of galectin biology beyond the predominant focus on LGALS3 and highlight LGALS1 as a potentially important transcriptional biomarker and candidate mediator of hypertension-related vascular stress in African Americans. Future longitudinal and mechanistic studies are warranted to determine whether galectin transcriptional signatures contribute directly to hypertension pathogenesis or reflect adaptive responses to chronic cardiovascular stress.

## Acknowledgements

This work was supported by the National Institute of Health under award numbers UC2MD019626, UC2GM162929, T32GM144927, and T32HL007737; by the United States (U.S.) Department of Veterans Affairs Office of Research and Development under award number 1I01CX002780; by the Chan Zuckerberg Initiative’s Foundation for Accelerate Precision Health Program to Advance Genomics Research at Meharry Medical College, CZIF2022-007043, and by the NIMHD grant U54MD007593 to build research capacity at Meharry Medical College. American Heart Association (https://doi.org/10.58275/AHA.25IVPHA1462290.pc.gr.229780 to MLL and GEG). Agencia Nacional de Promoción Científica y Tecnológica de Argentina (PICT 2019-02987 to GEG) and from Consejo Nacional de Investigaciones Científicas y Técnicas de Argentina (CONICET; PIP 938 to GEG).

## Perspectives

The galectin family has traditionally been viewed through the lens of LGALS3 and its role in cardiac inflammation and fibrosis in hypertension. However, our systematic transcriptomic evaluation reveals a much broader and heterogeneous network involved in cardiovascular homeostasis. The striking emergence of LGALS1 as the dominant transcriptional correlate of systolic blood pressure suggests that vascular stress responses may operate independently of the myocardial remodeling pathways predominantly governed by LGALS3. This paradigm shift implies that measuring individual galectins in isolation may obscure complex, context-dependent biological interactions. Instead, evaluating the relative balance within this network, such as the inverse association we observed with the LGALS3/LGALS1 ratio, could offer deeper insights into the inflammatory and homeostatic shifts that drive hypertensive pathogenesis. Future investigations should prioritize longitudinal and mechanistic studies to determine whether elevated LGALS1 expression is an early driver of blood pressure dysregulation or a compensatory defense mechanism against chronic hemodynamic stress. Expanding this multi-marker transcriptomic framework will be critical for unraveling the diverse functions of galectins and evaluating their potential as precision therapeutic targets in hypertensive cardiovascular disease.

## Novelty and Relevance

### What Is New?

- This study systematically compares four multiple galectin genes simultaneously within the same population, rather than evaluating them individually.
- LGALS1, emerged as the strongest transcriptomic correlate of systolic blood pressure in this population.

### What Is Relevant?

- The research highlights how distinct galectins relate to blood pressure in untreated African Americans, a population highly vulnerable to hypertensive complications.
- It demonstrates that galectins function as a diverse network with both positive and inverse effects on the cardiovascular system.

### Clinical/Pathophysiological Implications?

- The balance between different galectins (such as the LGALS3/LGALS1 ratio) may serve as a superior biomarker for vascular stress and immune-inflammatory shifts, paving the way for more precise hypertension monitoring and targeted interventions.

---

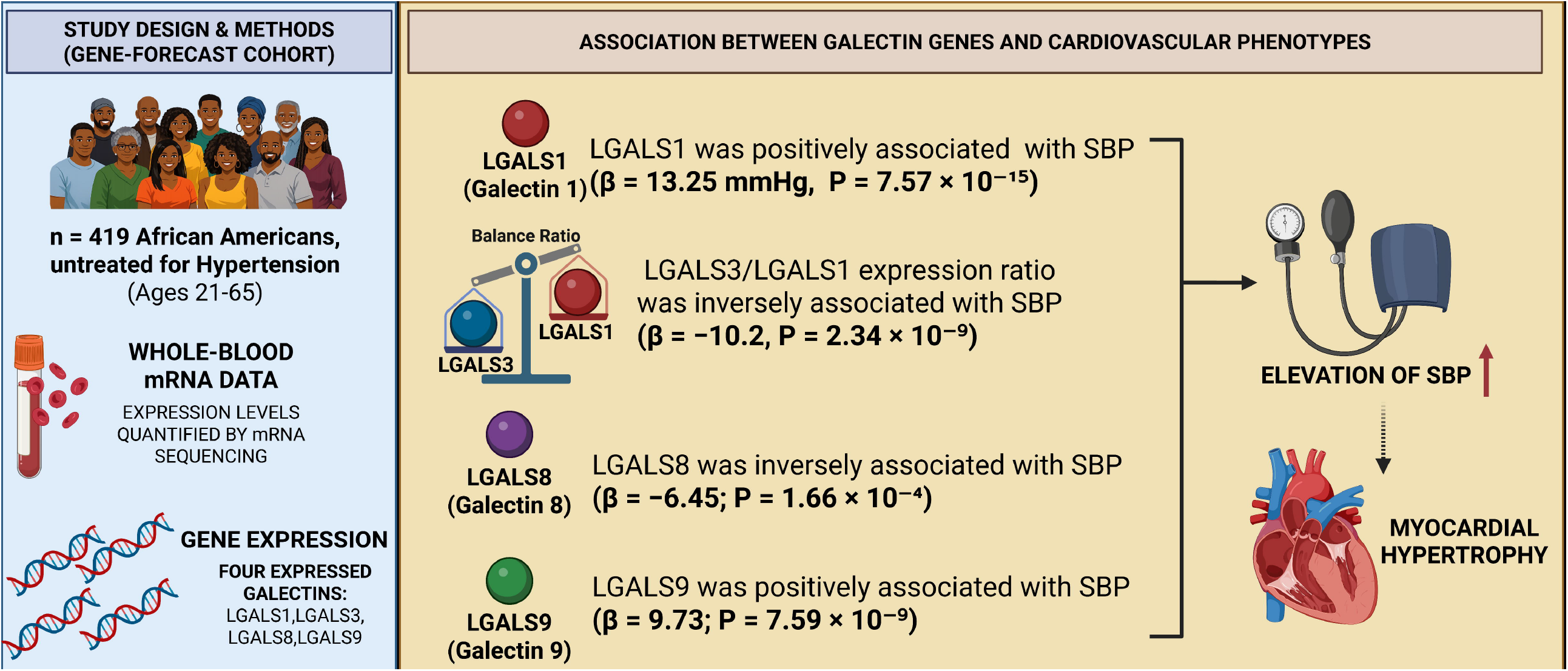

